# Associations Between Task-Related Modulation of Motor-Evoked Potentials and EEG Event-Related Desynchronization in Children with ADHD

**DOI:** 10.1101/2021.03.28.437305

**Authors:** Joshua B. Ewen, Nicolaas A. Puts, Stewart H. Mostofsky, Paul S. Horn, Donald L. Gilbert

## Abstract

Children with attention-deficit/hyperactivity disorder (ADHD) have previously shown a decreased magnitude of event-related desynchronization (ERD) during a finger-tapping task, with a large between-group effect. Because the neurobiology underlying several TMS measures have been studied in multiple contexts, we compared ERD and three TMS measures (Resting Motor Threshold [RMT], Short-Interval Cortical Inhibition [SICI] and Task-Related Up-Modulation [TRUM]) within 14 participants with ADHD (ages 8-12y) and 17 control children. The TD group showed a correlation between greater RMT and greater magnitude of alpha (10-13Hz, here) ERD, and there was no diagnostic interaction effect, consistent with a rudimentary model of greater needed energy input to stimulate movement. Similarly, inhibition measured by SICI was also greater in the TD group when the magnitude of movement-related ERD was higher; there was a miniscule diagnostic interaction effect. Finally, TRUM during a response-inhibition task showed an unanticipated pattern: in TD children, the greater TMS task modulation (TRUM) was associated with a smaller magnitude of ERD during finger-tapping. The ADHD group showed the opposite direction of association: greater TRUM was associated with larger-magnitude ERD. Prior EEG results have demonstrated specific alterations of task-related modulation of cortical physiology, and the current results provide a fulcrum for multimodal study.

---

Attention-Deficit/Hyperactivity Disorder (ADHD) is one of the most common neurobehavioral conditions in children, affecting around 4% of the pediatric population (Vasileva et al. 2020). Despite psychopharmacology and behavioral interventions that are effective in the short term in many cases, ADHD continues to have a substantial burden of negative outcomes (Molina et al. 2009), underscoring the need for novel diagnostics and therapeutics (Ewen 2016; Sahin et al. 2018).While diagnosis is based on the core, clinical symptoms of inattention, hyperactivity and impulsivity (Wolraich et al. 2019), cognitive research has repeatedly identified deficits in cognitive control, including response inhibition (Mostofsky and Simmonds 2008; Crosbie et al. 2013) as well as parallel deficits in motor control, particularly motor inhibition (Denckla and Rudel 1978). These specific deficits provide a mechanistic basis for the biology of ADHD for the purpose of developing new interventions and biomarkers to improve outcomes.

EEG is a technology that is of considerable interest for the development of biomarkers in a variety of conditions due to its relatively low cost and non-invasiveness. One particular EEG effect of interest for the development of biomarkers is event-related spectral perturbation, consisting of task-related increases (event-related synchronization; ERS) and decreases in oscillatory power (event-related desynchronization; ERD) (Pfurtscheller and Neuper 1994; Pfurtscheller and Lopes Da Silva 1999). Differences in ERS/ERD have been demonstrated in ADHD (McAuliffe et al. 2020) and other NDDs (Murphy et al. 2014; Ewen, Lakshmanan, et al. 2016). ERD refers to the relative suppression of a particular EEG oscillation during a cognitive or motor task. Our group has studied ERD in ADHD in the context of mirror overflow, a deficit in motor inhibition often found in children with ADHD (Cole et al. 2008; MacNeil et al. 2011; McAuliffe *et al.* 2020). Mirror overflow refers to the involuntary production of movement on the opposite side of the body from a volitional, unilateral movement.

We examined ERD in alpha (here, 10-13 Hz) and beta bands (here, 18-28 Hz). Both alpha and beta ERD are understood to be inhibitory in effect (Kelly et al. 2006; Engel and Fries 2010). The mu band (10-28 Hz), or sensory-motor rhythm, is an oscillation typically recorded from central scalp regions and is suppressed during a variety of motor tasks. Many studies report only the alpha component of the mu rhythm, but mu is truly composed of both alpha and beta components. Importantly these components show dissociations in experimental contexts (e.g., (McAuliffe *et al.* 2020)). Alpha is believed to be generated by the post-central gyrus (Salmelin et al. 1995) whereas beta activity is believed to be generated by pre-central gyrus (Keil et al. 2014). To set the stage for the current analysis, our prior results from within a larger sample, of which the sample presented here is a subset, demonstrated an ADHD-associated decrease in left-hemisphere alpha ERD, but not left-hemisphere beta ERD, during finger-sequencing of the right hand (eliciting mirror overflow in the left hand) (McAuliffe *et al.* 2020).

To maximize the utility of EEG as a tool for biomarker development, e.g., to index targets for treatment in clinical trials or other interventions, it is important to understand the mechanisms that underlie these EEG measures (Ewen and Beniczky 2018; Ewen et al. 2019; Ewen et al. 2021). While a good deal is known about the generators of mu-alpha and mu-beta rhythms and other cognitively-associated oscillatory activity (reviewed in (Cannon et al. 2014)), there is relatively little known about the mechanisms that allow for task-related modulation of them. Task-related modulation in particular is relevant, as baseline measures likely do not reflect the “real-life” implications of these cortical rhythms.

TMS, in contrast to EEG, has been studied in a wide range of demographic, diagnostic and pharmacological contrasts, and different TMS indices have been repeatedly demonstrated to reflect different aspects of physiology. For example, resting motor threshold (RMT) is the percentage of maximum stimulator output required to evoke a motor evoked potential (MEP) (Mills and Nithi 1997), such that a higher threshold indicates a greater energy requirement for activation and is thought of as a basic indicator of readiness of the motor cortex to depolarize. It may index terminal myelination, subcortical myelination and developmentally regulated changes in ion channels. A longstanding finding in healthy children is that RMT is highest in infancy and declines through childhood, reaching adult levels at approximately age 12 years (Muller et al. 1991).

Short-interval cortical inhibition (SICI) is one of the most widely studied TMS measures and utilizes paired-pulse approaches. SICI is quantified as the ratio of the conditioned (paired-pulse) MEP and unconditioned (single-pulse) MEP. SICI is understood to index GABA-A mediated inhibitory interneuronal activity acting on motor cortex (Kujirai et al. 1993). SICI is diminished in a large variety of neurodevelopmental and neurodegenerative disorders (Moll et al. 2001; Rothwell et al. 2009; Ni et al. 2013; Mimura et al. 2021). Altered SICI in ADHD is particularly robust, and correlates with parent-rated symptom severity of both hyperactive/impulsive and inattentive symptoms (Gilbert et al. 2011) and is modified by standard pharmacologic treatments (Moll et al. 2000).

The third TMS index reported was dubbed by our laboratory “TRUM”: Task-Related Up Modulation (of motor-evoked potentials) (Gilbert et al. 2019; Zea Vera et al. 2020). Interestingly, compared to rest, SICI diminishes during preparation to act and during actions (Garry and Thomson 2009; Hoegl et al. 2012; Gilbert *et al.* 2019) and is thus modulated by task-presence. Like the ERD, this metric is task associated. In the current set of experiments, TRUM was studied in the context of a modified, child-friendly version of the Slater Hammel Stop Signal Task (SST) (Guthrie Michael D. et al. 2018; Gilbert *et al.* 2019), which evaluates response inhibition performance. Different phases of the task are understood to represent action selection and response preparation. This research was based on prior observations that motor cortex excitability, and hence MEP amplitude, increases prior to cued and self-paced movements (Chen et al. 1998).

The goal of the current paper is to directly explore associations between task-based and resting measures of EEG and TMS indices. As many participants participated in only one procedure or the other, prior publications from larger samples of which the one presented here is a subset have reported independently on TMS (Gilbert *et al.* 2019) and EEG (McAuliffe *et al.* 2020). Within the larger sample that received TMS, we found that, compared with TD children, children with ADHD showed reduced SICI at rest and during the action-selection phase of the task (selecting to go or to stop), such that reduced SICI at rest does not resolve or normalize during engagement with this task. Additionally, we found that the amount of task-related up modulation (TRUM) of motor cortex excitability during this task was diminished in children with ADHD (Gilbert *et al.* 2019). However, SICI and TRUM were only minimally correlated with one another (Zea Vera *et al.* 2020), supporting the notion that these measures may be differentially sensitive to distinct mechanisms of altered physiology within ADHD.

The overall analytic strategy was first to examine EEG-TMS associations within the TD group alone, given the absence of pre-existing basic knowledge about the relationship between ERD and TMS indices. We next examined diagnostic (interaction) effects, looking for associations in which ADHD diagnosis may moderate the TMS-ERD relationship, as potentially promising areas of further study. Because of a dearth of TMS-EEG comparisons to date, our working hypotheses were necessarily speculative. Because only alpha ERD showed diagnostic group differences within the left hemisphere under the task conditions studied (the only hemisphere stimulated by TMS), we limited working hypotheses to those involving *alpha* ERD, though we explored beta ERD associations as well.

TD-only hypotheses were as follows: Because RMT reflects baseline “readiness to depolarize” (higher RMT = less readiness to depolarize), and ERD is analogous to the “energetic change” in the cortex, we predicted that TD children with higher RMT would show a greater magnitude of ERD. Our second prediction was that higher levels of SICI (thus, more suppression) would portend greater ERD, as a more inhibited resting cortex would require greater activation (indexed by ERD) to generate behavior. Thirdly, under the assumption that a joint “cortical physiology modulation” mechanism is indexed by TRUM and by ERD, we predicted that those two measures would correlate, despite the respective physiology being measured in the context of two separate tasks.

With regards to ADHD, within the larger TMS sample, RMT was not substantially decreased in the ADHD group (4.2% decrease in observed means compared with TD group, *p*=0.13) (Gilbert *et al.* 2019). Because alpha ERD was in fact different between groups, we predicted the presence of a diagnostic interaction effect. Secondly, the resting SICI was lower (i.e., ratio was increased) in the ADHD group (i.e., less inhibition) by 21% (*p*=0.03) (Gilbert *et al.* 2019), and alpha ERD was decreased (McAuliffe *et al.* 2020). We therefore expected SICI-alpha ERD correlations in both groups and no interaction effect. Finally, within the larger TMS sample, children with ADHD showed less TRUM (Gilbert *et al.* 2019). Within the larger ADHD sample, they showed less alpha ERD (McAuliffe *et al.* 2020). Therefore, we did not anticipate a group interaction effect within the combined sample (i.e., diagnosis does not moderate the TRUM-ERD relationship).

## MATERIALS AND METHODS

### Participants

Participants reflect a subsample of those reported by Gilbert *et al.*(Gilbert *et al.* 2019) and McAuliffe *et al.* (McAuliffe *et al.* 2020), each of which was a part of larger study with consistent recruitment criteria. Briefly, these were case-control studies of 8-12-year-old children with ADHD and typically developing (TD) controls. Children participated in the two studies within a 6-month time-period. ADHD diagnoses were based on parent interviews using the Diagnostic Interview for Children and Adolescents, Fourth Edition; (DICA-IV) (Reich 2000) or Kiddie Schedule for Affective Disorders and Schizophrenia for School-Aged Children (K-SADS) (Kaufman J. et al. 1997). TDs were excluded for any diagnosis on the DICA-IV/K-SADS or for elevated ADHD symptoms on the Conner’s Rating Scale-Revised (CPRS-R) (Conners et al. 1998). Additional exclusion criteria for both groups included history of seizures, intellectual disability, neurological illness or injury, or left-handedness/mixed dominance, as assessed by the Edinburgh Handedness Inventory (≤0.5) (Oldfield 1971). All children with full-scale IQ (FSIQ) scores below 80 on the Wechsler Intelligence Scale for Children, Fourth Edition (WISC-IV (Wechsler 2003) or WISC-V (Kaufman Alan S et al. 2015) were excluded.

### Finger-Tapping and EEG

#### Finger-Tapping Task used for EEG-ERD

Participants were instructed to tap each finger against the thumb in successive order (index-middle-ring-little) and self-paced timing, one hand at a time, for six seconds in a trial. A start cue was presented on a computer monitor. Left-handed finger-tapping (LHFT) and right-handed finger-tapping (RHFT) trials alternated in each block, although only RHFT trials were analyzed for this study. There were five blocks consisting of 20 trials in each per block. Behavioral overflow was measured in the non-tapping hand via electronic goniometers (Biopac Systems Inc., Goleta, CA); overflow was quantified per previous studies in our laboratory (MacNeil *et al.* 2011; McAuliffe *et al.* 2020).

#### EEG Recording, Preprocessing and Event-Related Desynchronization (ERD) Analysis

EEG was recorded during finger tapping using a 47-channel, full-scalp, equidistant WaveGuard cap system and an asa-lab amplifier (Advanced Neuro Technologies, Netherlands). Trials for each subject were excluded during a video analysis if children were observed not to be paying attention, moved out of compliance with visually displayed instructions, or did not complete at least 5 seconds of tapping within that trial. Data were recorded at a 1024 Hz sampling rate and 138 Hz anti-aliasing filter and were referenced to an average of all channels. Impedances were kept below 15 kΩ. Preprocessing was conducted in EEGLAB (Delorme and Makeig 2004). EEG data were preprocessed using asa-lab version 4 software. Data were high-pass filtered at 0.2 Hz, and visually inspected for eye-blinks, horizontal eye movements, and muscle activity. These artifacts could all be identified visually based on well-defined morphology. A Principal Component Analysis (PCA)-based method of removing artifact components within asa-lab was used to remove components that account for >90% of the variance of the artifact subspace. Not a single trial from any subject was removed in the artifact rejection step. To minimize effects of volume conduction, signals were then converted to current source density (CSD) estimates from CSD toolbox (Kayser and Tenke 2006) in MATLAB (Mathworks, Natick, MA). Full details can be found in McAuliffe *et al.* (McAuliffe *et al.* 2020).

The EEG measures of interest were alpha band (empirically derived 10-13 Hz range) and beta band (empirically derived 18-28 Hz range) ERD in the scalp region approximating left M1, during right hand finger tapping (RHFT). Bands were selected via spectrogram, as reported in (McAuliffe *et al.* 2020). Analysis was restricted to Left M1/RHFT because the TMS procedures only interrogated left M1 via EMG captured from right hand. Data were down-sampled to 256 Hz, and ERD was calculated for each channel as follows: at each time-frequency point during the task (starting from the point of tapping onset for each trial, as measured by initial goniometer deflection in the tapping hand), a *z*-score was calculated relative to a distribution created from the baseline period (1s prior to start cue). We limited our analysis to a 1.5 second window (1.5– 3s relative to tapping onset) in the middle of each tapping block to avoid EEG onset and offset (rebound) effects. ERD-related *z*-scores for each channel were integrated over 384 time-samples (in 1.5 seconds) × 8 frequency bins per Hz. ERD is a negative value, so a greater magnitude of ERD is a more negative value.

### Stop-Signal Task and TMS

#### Slater Hammel Stop-Signal Task (SST)

Participants operated a standard game controller with their right, dominant hand for this response inhibition task. GO and STOP Stimuli were presented on a computer monitor via Presentation (v.10.0; Neurobehavioral Systems, Albany, CA). Ulnar aspects of both arms and hands rested on a body-surrounding pillow (The Boppy Company, LLC, Golden, CO) so the palmar surface faced medially. The dominant hand operated the game controller with a fully extended index finger. Surface EMG electrodes recorded the first dorsal interosseous (FDI) muscle. The participant initiated each trial by adducting (pushing down) the index finger on the game controller button, activating the finger flexors (antagonistic to the FDI), causing a racecar at the left side of the screen to audibly start its “engine” and then traverse a straight, 1000 ms “racetrack” across the screen. The car kept going only as long as the finger is adducted. The “go action” of this task required lifting the finger, i.e., activating FDI, when the car was as close as possible to the 800 ms mark, without going past it. However, in 25% of trials, at random, the car stopped itself spontaneously 300 to 700 ms after trial onset. This was the “Stop Cue.” The child was instructed that if the car stops itself early, they should suppress their finger lift action and maintain their finger pressed down until they saw a checkered flag (which occurs at 1000 ms). Successful stopping is “not lifting the finger at the 800 ms mark,” and maintaining finger adduction for greater than 1000 ms. The stop cue timing shifted by 50 ms increments depending on success or failure, allowing the stop trial times to converge to indicate response inhibition efficiency (Coxon et al. 2006; Guthrie Michael D. *et al.* 2018). The game was played with three 40 trial blocks (30 go with 10 stop randomly intermixed). A full description and demonstration of the task is available open-access (Guthrie Michael D. *et al.* 2018).

#### TMS in Resting Motor Cortex (RMT, SICI)

Dominant (left) hemisphere M1 physiology was assessed using a Magstim 200^®^ transcranial magnetic stimulators (TMS) (Magstim Co., New York, NY, USA) connected through a Bistim^®^ module to a round 90 mm coil and Signal^®^ processing software as described previously (Gilbert et al. 2011; Guthrie Michael D. *et al.* 2018). TMS utilizes magnetic fields to generate an electric field that can induce depolarization in neurons within range of the coil. Single suprathreshold intensity pulses over M1 can generate a motor evoked potential (MEP) measurable in anatomically localized muscles with surface EMG. Pairing suprathreshold pulses with preceding subthreshold TMS pulses can consistently inhibit or activate motor cortex interneurons, reducing or increasing the amplitude of the MEP. TMS coil placement was flat at the vertex, with the handle directly posterior. This technique and coil were chosen to enhance stability in hyperkinetic children (compared to the more common tangential placement of the figure of 8 coil). All protocols for active and resting motor thresholds (AMT, RMT) (Mills and Nithi 1997) and paired-pulse TMS for SICI (Kujirai *et al.* 1993; Rothwell *et al.* 2009) are in standard use, implemented by our laboratories in 8 to 12 year old children, as previously described (Gilbert *et al.* 2011). In brief, threshold measures were performed first, to habituate children, starting with pulses at 10% maximal stimulator output, increasing by 10% until a consistent MEP was observed, then decreasing the intensity until a minimum point was reached where 3 of 6 pulses produced no MEP and 3 produced an MEP of approximately at least 50 microvolts, at rest (RMT). RMT was indexed by the maximum stimulator output of the TMS stimulator being used; the maximum is 100%.

SICI at rest was evaluated using 10 single test pulses and ten paired pulses, with conditioning pulses at 0.6×RMT and test pulses at 1.2×RMT, at an interstimulus interval of 3 ms and an inter-trial interval of 6 s, ±5%. Test pulses were administered at 1.2×RMT with a goal of evoking MEPs averaging 0.5 to 1.5 mV per trial. SICI is expressed as ratios of paired to single pulses. For SICI, ratios closer to 1.0 indicate less inhibition by the 3 ms pair, relative to the single pulse, i.e., less SICI. 17 TD participants and 14 with ADHD underwent SICI testing.

#### TRUM

TRUM was measured during the SST; there were two blocks of 40 trials. Single (at 1.2×RMT) and 3-ms-paired (at 0.6 and 1.2×RMT) TMS pulses were delivered randomly across trials. For go trials, TMS pulses were administered at the time of expected action selection – 150 ms prior to the finger lift; for stop trials, TMS pulses were administered at the time of expected action suppression – not lifting the finger, 150 ms after the stop cue. TRUM is the ratio of MEP amplitudes during the response inhibition task compared to MEP amplitudes at rest. 8 TD participants and 9 with ADHD underwent TRUM testing.

### Statistical Approach

This is a post-hoc, exploratory analysis from an intersecting sub-sample of previously published data from two unique studies involving analysis of well-characterized children with ADHD and TD controls (Gilbert *et al.* 2019; McAuliffe *et al.* 2020). Diagnostic group comparisons were performed with unpaired *t-*tests for continuous variables and χ^2^ for categorical variables. We calculated group distributional effect sizes as Cohen’s *d,* as measures with the greatest diagnostic-group separation would likely be the most effective diagnostic biomarkers (Group 2016; Ewen *et al.* 2019).

The relationship between RMT and ERD (alpha and beta, separately) was tested using linear models separately in each group, adjusting for sex, age (which is known to affect RMT) and general intellectual ability (General Ability Index of the Wechsler tests: GAI), which showed nearly significant differences between groups.

The point estimate of SICI was calculated as the average of paired pulse TMS-evoked MEP amplitudes divided by average of single pulse TMS-evoked MEP amplitudes. SICI-ERD associations were tested using a repeated measures mixed models with MEP amplitude as the dependent variable, ERD as a predictor variable, and subject as a random effect, incorporating all trials instead of averaging within subjects (Guthrie M. D. et al. 2018; Gilbert *et al.* 2019). Age, sex and GAI were also included in the models. Log transformation was used to optimize residuals. In repeated measures, SICI was estimated from the PulseType (paired vs. single), and the SICI-ERD association as PulseType×ERD. Rather than stratifying, diagnostic effects were then estimated from interaction terms Diagnosis×PulseType, where Diagnosis was a categorical term. The test for diagnostic difference was thus a three-way interaction term: MEP = Diagnosis×PulseType×ERD.

TRUM was calculated using analogous models, except that the TRUM effect was defined by the Block (task *vs*. rest) variable, the TRUM-ERD association as Block×ERD, and the test for diagnostic difference as the significance of the three-way term: MEP = Diagnosis×Block×ERD.

*P* values less than 0.05 were considered significant, and no correction was made for multiple comparisons. All models were analyzed using SAS^®^ statistical software version 9.4 (SAS Institute Inc., Cary, NC). Relationships with ERD were determined by including alpha and beta ERD as factors in separate models.

## RESULTS

### Participants

There were 17 children in the TD group (75% male; age 10.9±1.4; 10.2-11.6 years; 59% Caucasian, 6% African American, 18% Asian, 18% Biracial, 0% Hispanic ethnicity) and 14 in the ADHD group (57% male; age 10.7±1.2; 10.1-11.4 years; 71% Caucasian, 7% African American, 0% Asian, 0%, 21% Biracial, 14% Hispanic ethnicity) who had both EEG and TMS data available (see Table 1).

**Table 1.** Participants

|  | ADHD |  |  | TD Control |  |  | p-values |
| --- | --- | --- | --- | --- | --- | --- | --- |
|  | n | Mean | SD | n | Mean | SD | t-test |
| **<u>Age</u>** | 14 | 10.7 | 1.2 | 17 | 10.9 | 1.4 | 0.74 |
| **<u>ADHD severity</u>** |  |  |  |  |  |  |  |
| Conners T score Hyper/Impulsive | 14 | 75 | 15 | 17 | 45 | 6.8 | <.0001 |
| Conners T score Inattentive | 14 | 75 | 11 | 17 | 45 | 9.8 | <.0001 |
| ADHD-RS Hyper/Impulsive | 13 | 11 | 7.6 | 17 | 2.0 | 2.4 | 0.0005 |
| ADHD-RS Inattentive | 13 | 18 | 4.6 | 17 | 2.7 | 2.9 | <.0001 |
| ADHD-RS Total | 13 | 30 | 10 | 17 | 4.8 | 4.8 | <.0001 |
| **<u>Cognitive</u>** |  |  |  |  |  |  |  |
| Full Scale IQ | 13 | 107 | 10 | 13 | 116 | 12 | 0.047 |
| GAI | 14 | 104 | 13 | 17 | 112 | 13 | 0.081 |
| **<u>Other</u>** |  |  |  |  |  |  |  |
| Hollingshead Socio-economic | 12 | 54 | 11 | 16 | 59 | 6 | 0.18 |

ADHD attention deficit hyperactivity disorder. ADHD-RS rating scale. IQ intelligence quotient. GAI general intellectual ability. TD typically developing.

### Between-Group Effect Sizes

Of the various physiological metrics, alpha ERD during finger-tapping showed the greatest between-group separation (Cohen’s *d*=0.89, “large” effect, by convention). RMT showed *d*=0.5, and TRUM during the SST *d*=0.42, both “medium” effect sizes. Baseline SICI and beta ERD both showed effect sizes of 0.35 and 0.31, respectively, both in the “small to medium” range.

### RMT and ERD

Consistent with our working hypothesis, children with higher resting thresholds had greater ERD in the alpha range, after adjusting for age, sex and GAI (see figure 1, reporting RMT residualized for age, sex and GAI). This association was statistically significant in the TD group (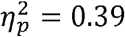; *p*=0.024; one outlier removed). While it did not quite reach significance in children with ADHD, the measured effect size was similar (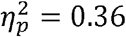; *p=*0.068; one outlier removed), and there was no significant diagnostic interaction effect (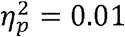; *p*=0.62), contrary to our predictions (Fig. 1). The association between RMT and beta ERD was marginal at best in TDs (; *p=*0.14; one outlier removed) and non-significant in the ADHD group (; *p=*0.97; one outlier removed). There was no significant diagnostic-group interaction (; *p*=0.65) (Fig. 2).

**Figure 1.**
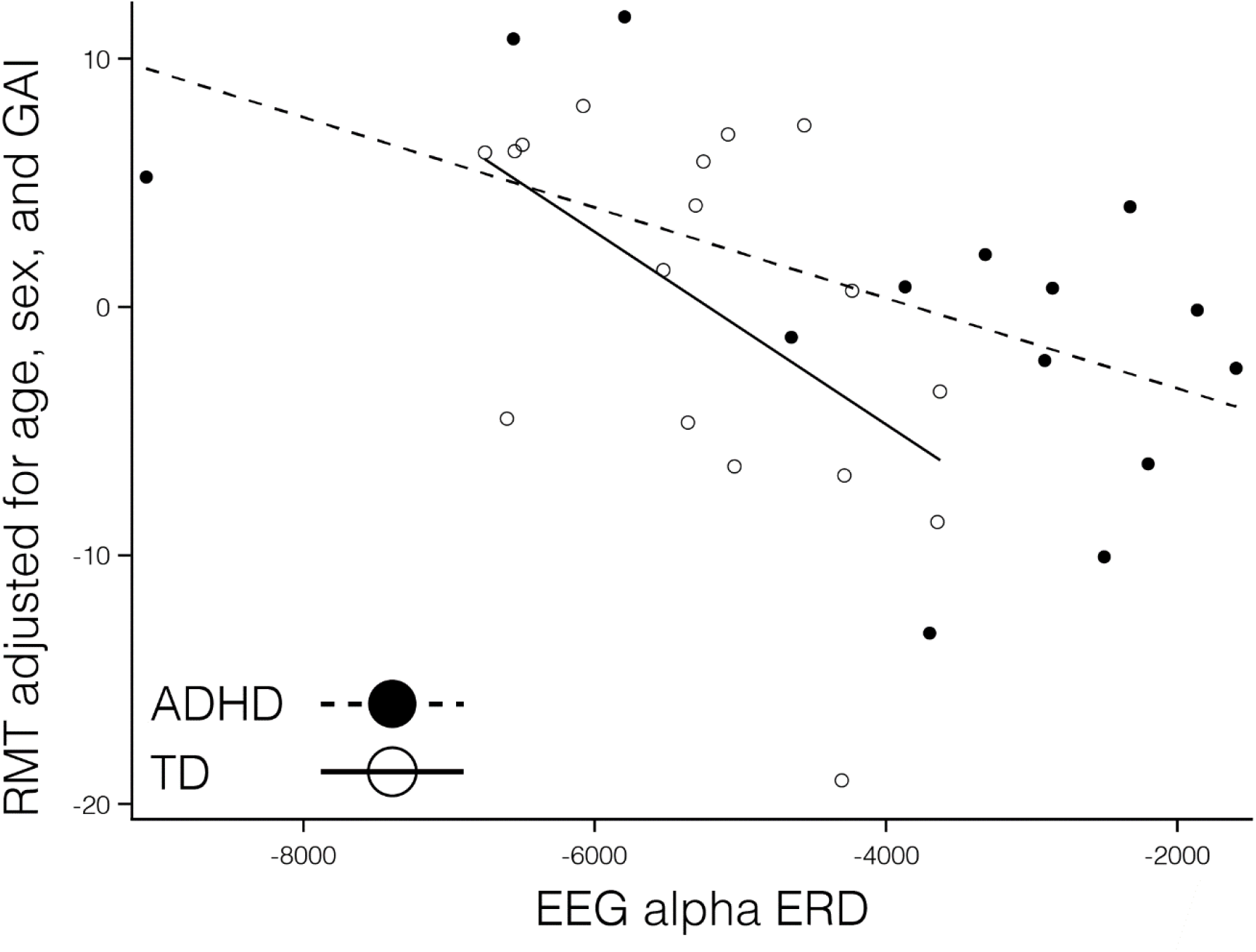
There was a significant association between RMT (adjusted) and alpha ERD in the TD children (hollow circles and solid line; *p*=0.024; with one highly influential point removed). The association in the ADHD group did not reach significance though the measured effect size was similar (; *p*=0.068; with one highly influential point removed). There was no significant diagnostic interaction effect (*p*=0.62), contrary to our predictions.

**Figure 2.**
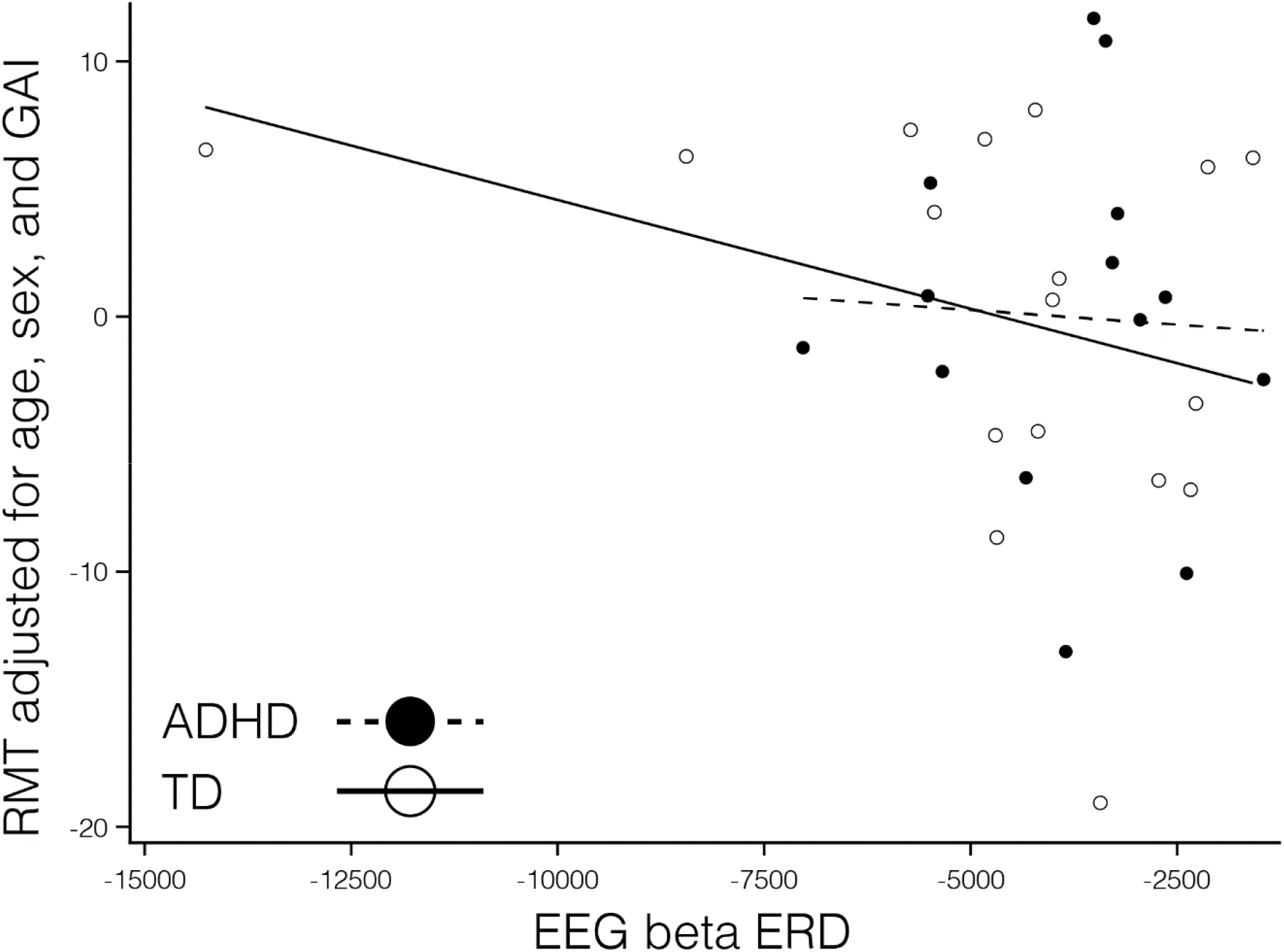
There was no statistically-significant association between RMT and beta ERD in TDs (; *p=*0.14; one influential point removed) or in the ADHD group (; *p=*0.97; one influential point removed). There was no significant diagnostic-group interaction (; *p*=0.65). Note that the outlier to the negative end of the *x-*axis was not influential; removing it did not substantially alter the calculated statistical results.

#### Short Interval Cortical Inhibition (SICI) and Event Related Desynchronization (ERD)Alpha ERD

Baseline SICI was significantly associated with alpha ERD in the TD control group, after adjusting for age, sex and GAI (*p*=0.0024, *n*=17), with a greater magnitude of ERD corresponding to a greater SICI ratio (interpreted as greater inhibition). SICI was not associated with alpha ERD in the ADHD group (*p*=0.27, *n*=14). While the diagnostic interaction effect reached statistical significance (*p*=0.017), the estimated effect size was small (Fig. 3). An approximation of for this repeated-measures model based on the *F-*statistic divided by the quantity (*F*-statistic + degrees-of-freedom of the denominator) came to a value of 0.01.

**Figure 3.**
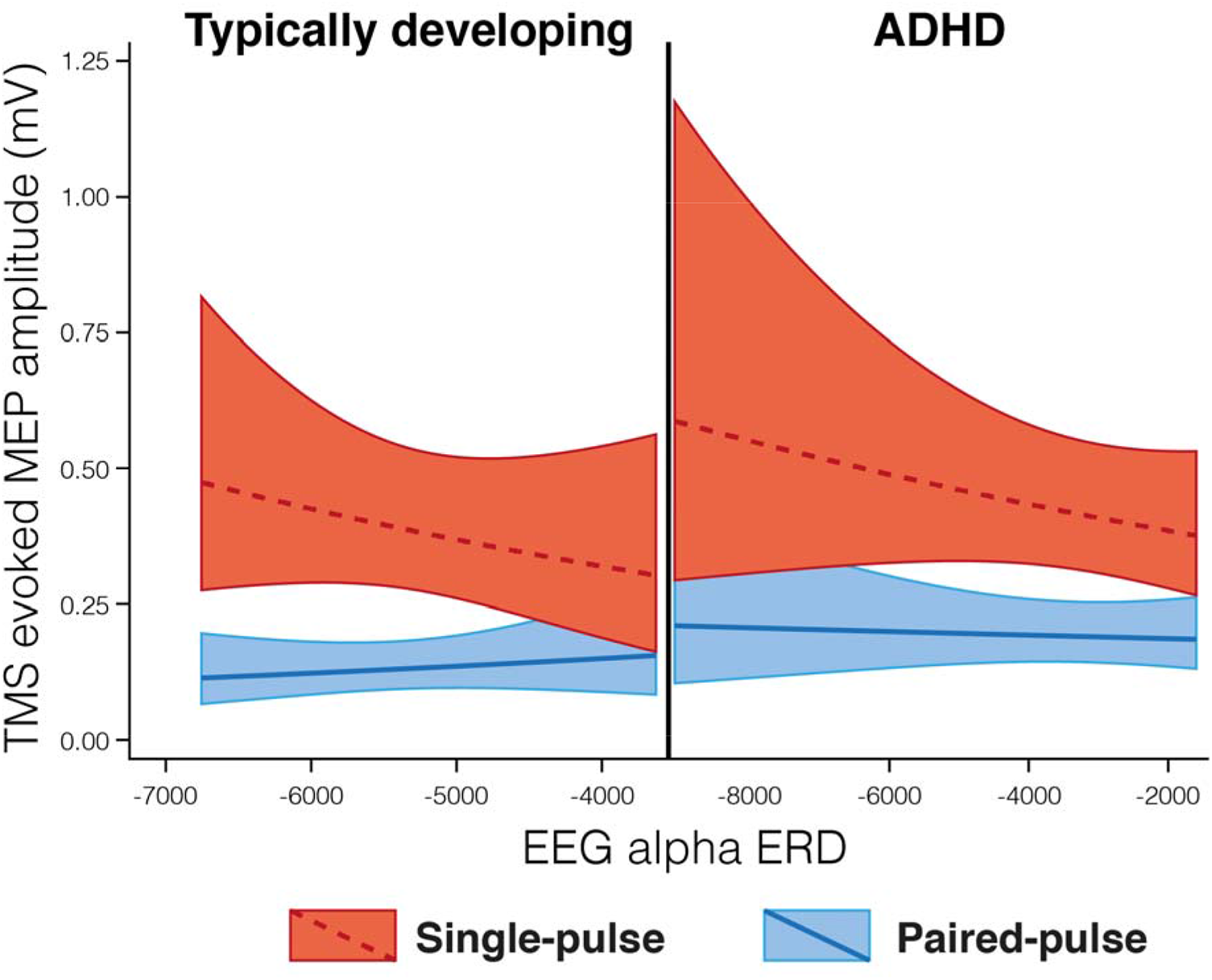
Regression Mixed-Model alpha ERD/baseline SICI relationships in *n*=17 typically developing (TD) children and *n*=14 children with ADHD. In these figures, the upper (dashed) line is single pulse MEP amplitudes, the lower (solid) line is 3 ms paired (inhibitory) MEP amplitudes. A greater distance between the two lines indicates a greater SICI ratio (i.e., more inhibition), and a more negative ERD value indicates a higher magnitude of ERD. The baseline SICI-alpha ERD association reached statistical significance in the control group (*p*=0.0024) but not in the ADHD group (*p*=0.27), with a statistically significant but very-small-magnitude diagnostic interaction effect (*p*=0.017).

#### Beta ERD

We found no evidence of an association between magnitude of beta ERD and baseline SICI in the TD group after adjusting for age, sex and GAI (*p*=0.85), however there was a significant association in the ADHD group (*p*=0.045) (Fig. 4), with a greater magnitude of ERD corresponding to a greater SICI ratio (interpreted as greater inhibition). There was no significant diagnostic-group interaction effect (*p*=0.11), though the groups did show different directions of observed effect, with the TD group showing a smaller SICI ratio associated with a greater magnitude of ERD (Fig. 4).

**Figure 4.**
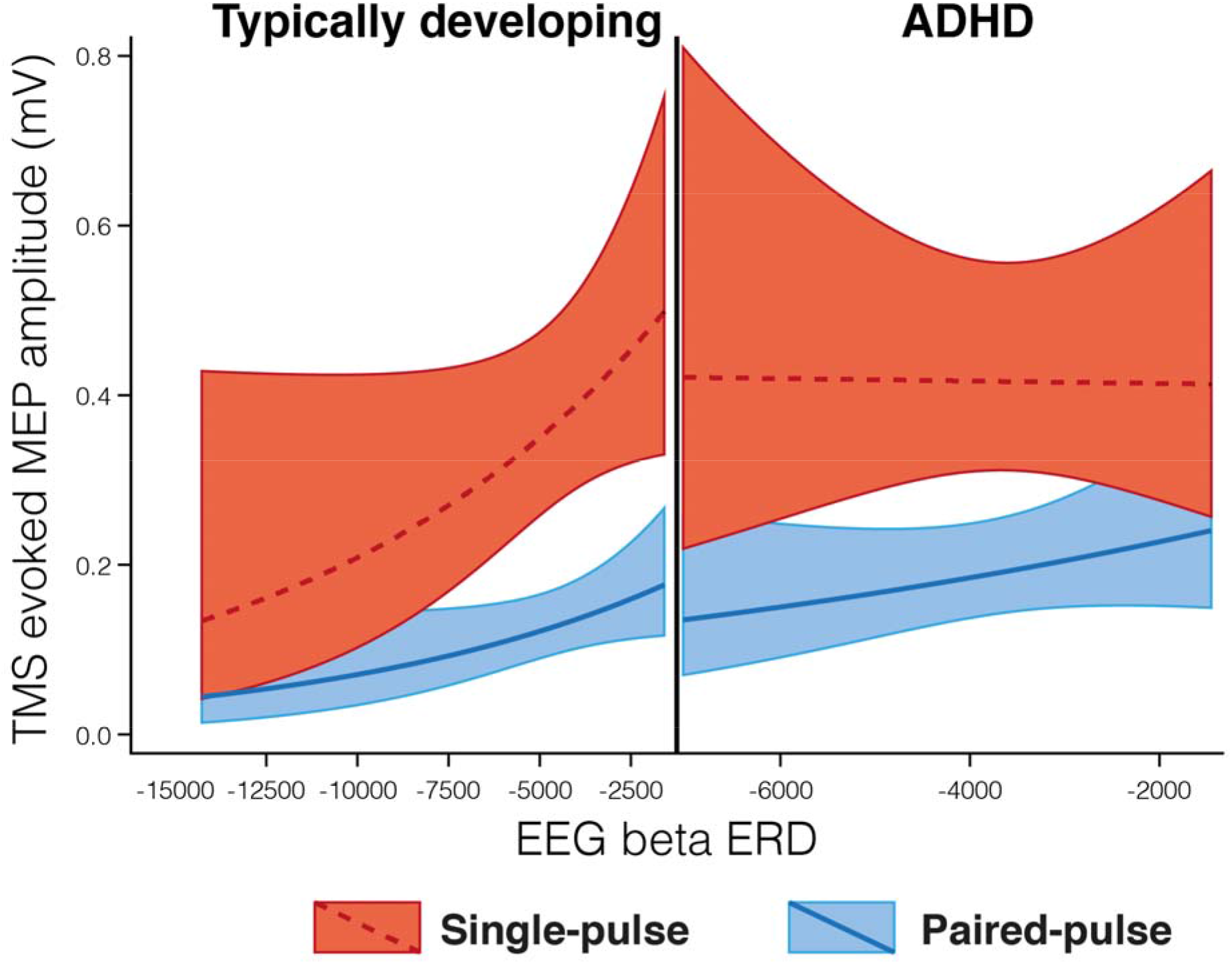
Regression Mixed-Model beta ERD/SICI relationships in *n*=17 typically developing (TD) children and *n*=14 children with ADHD. In these figures, the upper (dashed) line is single pulse MEP amplitudes, the lower (solid) line is 3 ms paired (inhibitory) MEP amplitudes, with a greater distance between the two lines indicating a larger SICI effect. A more negative ERD value indicates a greater magnitude of ERD. There was no statistical association between SICI and beta ERD in the TD group (*p*=0.85), however there was a significant association in the ADHD group (*p*=0.045). There was no significant diagnostic-group interaction effect (*p*=0.11), though the plot illustrates an opposite direction of association between groups, with TD showing a smaller SICI ratio associated with a larger magnitude of ERD and ADHD showing a larger SICI ratio associated with a larger magnitude of ERD. The non-linear shape in TD is due to logarithmic transformation to optimize residuals.

#### Task-Related Up Modulation (TRUM) and Event Related Desynchronization (ERD)Alpha

After removing one outlier (TD group), higher alpha ERD in left M1 during right-hand finger tapping was associated with *lower* magnitude of TRUM (*p*<0.001) after adjusting for age, sex and GAI, i.e., in the direction opposite from our hypotheses. By contrast, for children with ADHD, higher alpha ERD was associated with a *greater* magnitude of TRUM (opposite direction to TD group) (*p*<0.001). There was a large diagnostic interaction effect (*p*<0.001), contrary to our predictions (Fig. 5).

**Figure 5.**
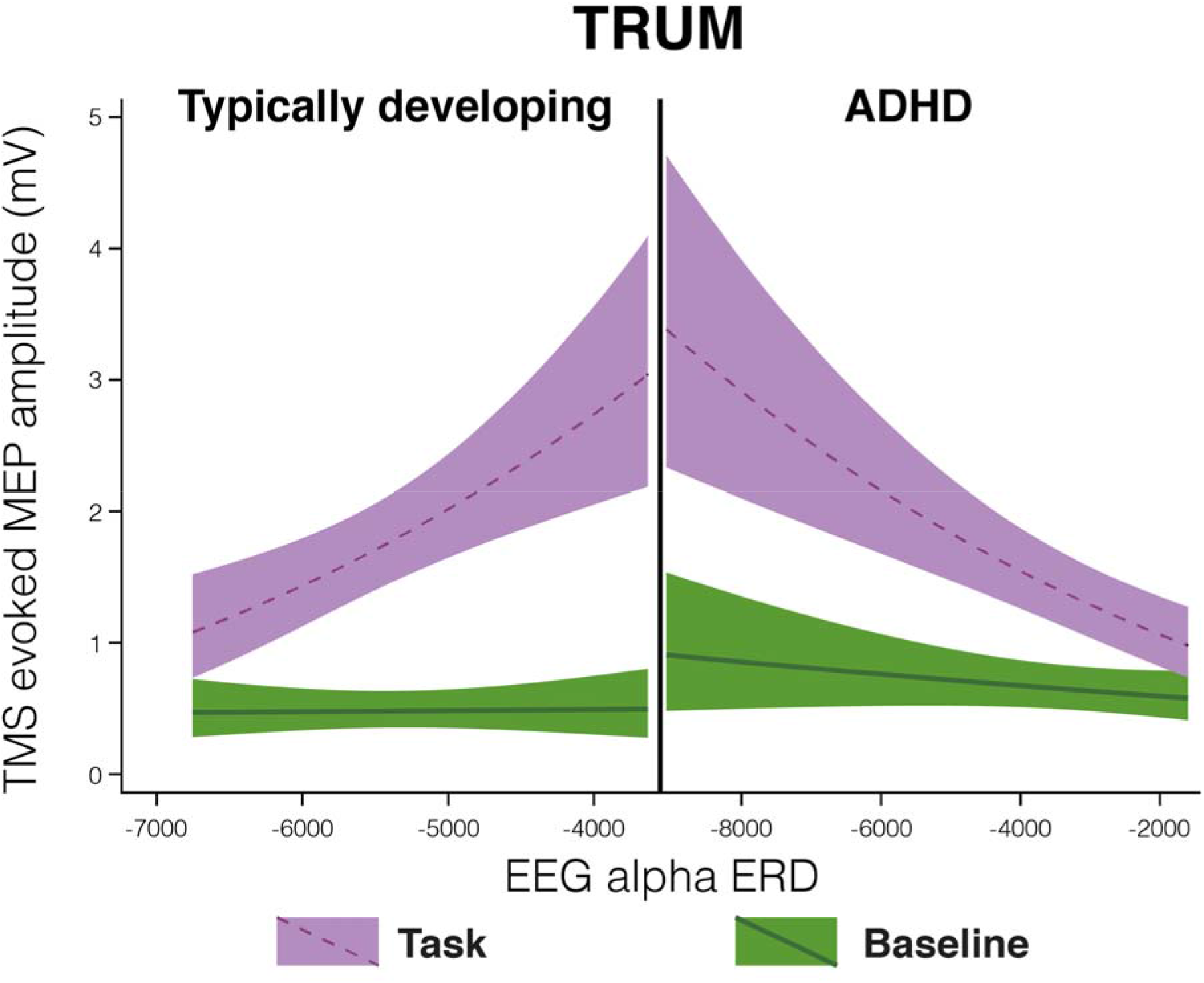
Regression Mixed-Model alpha ERD/TRUM relationships in *n*=8 typically developing (TD) children and *n*=9 children with ADHD. In these figures, the upper (red) line is MEP amplitudes during the response inhibition task trials, the lower (blue) line is the MEP amplitudes at rest. Among TD children with more left M1 alpha ERD when finger tapping, there is less left M1 TRUM (the ratio of the red to the blue line value) (*p*<0.001). The relationship is opposite children with ADHD (*p*<0.001), and the diagnostic interaction effect was large (*p*<0.001). The non-linear shape is due to logarithmic transformation to optimize residuals.

#### Beta: Left M1

In TD children, there was a statistical association between TRUM and beta ERD (*p*<0.001), with greater TRUM associated with a *decreased* magnitude of ERD, as also seen in the *alpha* ERD-TRUM relationship. In the ADHD group, the relationship between TRUM and beta ERD was again similar to that seen with alpha ERD, with a greater magnitude of TRUM associated with a greater magnitude of ERD (*p*=0.021). There was a large TRUM-beta ERD diagnostic interaction effect (p<0.001) (Fig. 6).

**Figure 6.**
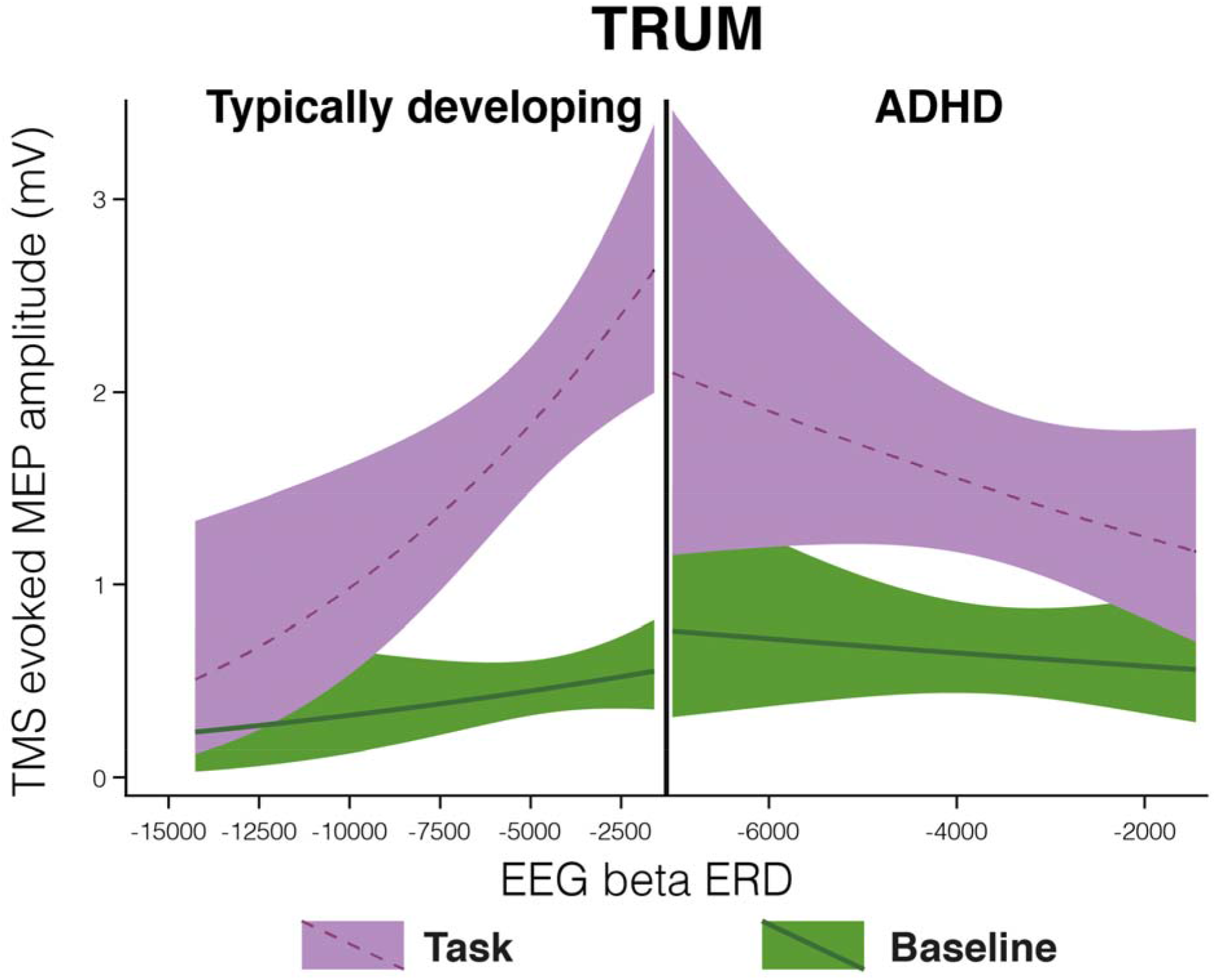
Regression Mixed-Model beta ERD/TRUM relationships in *n*=8 typically developing (TD) children and *n*=9 children with ADHD. In these figures, the upper (red) line is MEP amplitudes during the response inhibition task trials, the lower (blue) line is the MEP amplitudes at rest. The pattern of results was similar to that seen with alpha ERD: i.e., TD children showed a greater magnitude of ERD when the magnitude of TRUM was smallest (*p*<0.001), and the children with ADHD showed a greater magnitude of ERD when the magnitude of TRUM was *largest* (*p*=0.021). The was a diagnostic interaction effect (*p*<0.001). The non-linear shape in the TD group is due to logarithmic transformation to optimize residuals.

## DISCUSSION

Our primary findings in the TD-only analyses were consistent with two of three of our working hypotheses for that population group: RMT correlated with both alpha and beta ERD, such that a higher TMS field strength needed to generate a MEP was associated with a greater magnitude of ERD during the finger-tapping task. Similarly, SICI and alpha ERD correlated, such that greater SICI (i.e., greater inhibition) at rest was associated with a greater magnitude of alpha ERD. And while TRUM was statistically associated with both alpha and beta ERD, the direction was opposite to the one predicted for TD children: greater task-related up-modulation of MEP from rest to task engagement was associated with a *smaller* magnitude of alpha ERD during the finger-tapping task.

Regarding effects of diagnosis (ADHD vs. TD), for our primary RMT-alpha ERD finding, there was no significant diagnosis×physiology interaction in either the alpha or beta band, with significant positive associations in both ADHD and TD children. For SICI, there was also no evidence of an interaction effect (though ADHD *post-hoc* testing showed no evidence of a SICI-alpha ERD association; the presence of a statistical association in TD but absence in ADHD could be due to greater heterogeneity in the ADHD group). The TRUM-ERD×diagnosis effect was pronounced, and the association was unexpectedly in the opposite direction for the two groups, with greater task-related up-modulation of MEP from rest to task engagement associated with a *larger* magnitude of both alpha and beta ERD during the finger-tapping task.

### RMT, SICI and ERD

Our working model relating RMT and ERD assumed that RMT and ERD reflected aspects of a single mechanism and suggested that a larger RMT would indicate less “readiness to depolarize,” and a greater magnitude of ERD would be required to activate the cortex. The results are consistent with this hypothesis. However, the results presented here further assumed that all participants “reach the same point” of activation during finger tapping. It may be warranted, in future work using larger samples, to use additional task-related physiological measurements or behavioral measurements to quantify the degree of task-related activation on an individual-participant or even trial-by-trial basis. Such a model would test the notion that RMT+ERD=degree of task-related activation.

The SICI-ERD results also showed an association with alpha ERD, but only in the TD group. As SICI is believed to be mediated by GABA-ergic interneuron input into motor cortex pyramidal cell output, this suggests that the nature of SICI/ERD interactions might be clarified in future multi-modal studies in which participants also undergo GABA measurements with MRS (Harris et al. 2021 (in press)). However, given the lack of association of RMT and SICI in the TMS supra-sample (Gilbert *et al.* 2019), this suggests that alpha ERD magnitude is likely dependent on two separate mechanisms, one indexed by RMT and the other by SICI. Failure to find this SICI-ERD relationship in ADHD is difficult to interpret but may related to heterogeneity of ADHD as a categorical diagnosis reducing the signal to noise ratio.

### TRUM and ERD

The most robust and strikingly unexpected findings in this dataset was highly significant diagnostic interaction effect, such that:1) opposite to what we predicted, for TD children, higher alpha ERD in left M1 during right-hand finger tapping was associated with *lower* magnitude of TRUM; yet, 2) consistent with what we predicted for the TD group, for children with ADHD, higher alpha ERD was associated with a *greater* magnitude of TRUM. There are a few conclusions that we can derive from these results. Firstly, TRUM and ERD do not reflect the same underlying process, as we initially assumed. This is borne out by the anti-correlation within the TD group as well as the diagnosis-related interaction effect. However, the presence of TRUM-ERD statistical associations demonstrates that these processes interact. What these processes are, how they interact and how dependent they may be on the difference in cognitive-motor task between the TMS (SST) experiment and the EEG (finger-tapping) experiment needs to be explored with additional data and additional tasks.

The literature linking task-related modulation of TMS and EEG (i.e., TRUM and ERD) is sparse. LePage *et al.* (Lepage et al. 2008) systematically examined the dissociation between task-related TMS modulation and *alpha* ERD results in 16 healthy adults, albeit during a different series of motor tasks (as opposed to our cognitive control task), finding, as did we, a lack of correlation between TMS modulation indices and alpha ERD. Given strong evidence for reliability and validity of each measure, and assuming that TMS modulation and ERD reflect the same processes, they attributed the lack of correlation to the proposition that alpha ERD is more reflective of post-central activation, whereas beta (which they had not measured) was reflective of pre-central activation. They predicted that future work examining beta ERD (reflective of the physiology of the precentral gyrus) would indeed find correlations with task-related TMS modulation. Our results belie that hypothesis. Taken together, both our results suggest that TMS measures of modulation and ERD are sensitive to different task-engaged processes.

What is clear in our dataset is that the largest between-group effect size comes from strong diagnosis-associated effects associated with comparing *task-related modulation* of both EEG and TMS. As noted in the Introduction, the phenomenon of altered task-related modulation of brain physiology has been identified within not only ADHD (McAuliffe *et al.* 2020) but also autism spectrum disorder (ASD) (Murphy *et al.* 2014; Ewen, Pillai, et al. 2016; Pillai et al. 2017; Harvy et al. 2019), even in instances where baseline measures did not differ (or differ much) between groups. While our basic neuroscientific understanding is strong in relation to many aspects of resting brain physiology, including the TMS measures covered above as well as the generation of EEG oscillations (Cannon *et al.* 2014), the associated understanding of mechanisms of task-related modulation of brain physiology is limited. The strong diagnosis-specific signal from the TRUM-ERD analyses appears to support mechanisms of cortical modulation as a promising area of targeted investigation, particularly as it relates to NDD, and could lead to novel biomarkers and treatments. Trans-diagnostic research is key to demonstrating what aspects of these various modulatory effects are task-specific (as also in (Lepage *et al.* 2008)), diagnosis-related or related to the co-morbidity among these conditions.

## Acknowledgements

We appreciate the generous involvement of our research participants and their families as well as the laboratory members who were involved in data collection and analysis.

